# A tool for computation of changes in Na^+^, K^+^, Cl^−^ channels and transporters due to apoptosis by data on cell ion and water content alteration

**DOI:** 10.1101/486811

**Authors:** Valentina E. Yurinskaya, Igor A. Vereninov, Alexey A. Vereninov

## Abstract

The study aims to know how the apoptotic alteration of cell ionic balance follows from the quantitatively characterised time dependent decrease in the sodium pump rate constant and changes in permeability coefficients of Cl^−^, K^+^, and Na+ channels. New experimental data on changes in cell K^+^, Na^+^, Cl^−^, water contents, and the Na^+^/ K^+^-ATPase-mediated K^+^ influx during the first 4 h of the staurosporine (STS) induced apoptosis are used as a basis for quantitative characterisation of channels and transporters responsible for apoptotic cell ion balance alteration. New computational tool is developed. It is found that the dynamics of alteration of ion and water balance in the studied U937 cells were associated with the decrease in the Na^+^/K^+^-ATPase rate coefficient by 2.2 times for 4 h, and a time-dependent increase in potassium channel permeabilitry, and a decrease in the sodium channel permeability, whereas the early decrease in [Cl^−^]_i_ and cell volume were associated with an approximately 5-fold increase in the chloride channel permeability. The developed approach and the provided executable file can be used to identify the channels and transporters responsible for alterations of cell ion and water balance not only during apoptosis but in other physiological scenarios.

## Introduction

A characteristic feature of apoptosis, one of the basic genetically encoded cell death mechanisms, in contrast to accidental death, is that it is not associated with cell swelling or plasma membrane rupture (Galluzzi et al. 2018). The apoptotic volume decrease (AVD) in cells is a common although unnecessary hallmark of apoptosis (Maeno et al. 2000, 2006; Okada et al. 2001; Yurinskaya et al. 2005*a*, *b*; Bortner & Cidlowski, 2007). Cell swelling in apoptosis is prevented by the specific alteration of the monovalent ion balance in apoptotic cells, as the monovalent ions are major cell water regulators. It is believed that monovalent ions play an important role in apoptosis (Lang et al. 2005; Lang and Hoffman, 2012; Bortner and Cidlowski, 2014; Hoffmann et al. 2015; Kondratsky et al. 2015; Jentsch, 2016; Wanitchakool et al. 2016). However, this opinion is based mostly on the fact that ion channels and transporters are altered somehow during apoptosis and that their pharmacological or genetic modification has an effect on apoptosis. The mechanism of specific apoptotic alteration of cell ion and water balance has gotten much less attention than the molecular identity of channels and transporters involved in apoptosis. The mechanistic studies are hampered by the interdependence between ion fluxes via the numerous channels and transporters in the plasma membrane. This difficulty can be overcome by the computational analysis of whole-cell ion flux balance, which has been developed for normal cells (Jakobsson, 1980; Lew and Bookchin, 1986; Lew et al. 1991; Terashima et al. 2006; Vereninov et al. 2014, 2016). However, no successful analyses have been done on apoptotic cells. We have studied the relationships between alterations of the sodium pump or K^+^, Na^+^, and Cl^−^ channels and transporters and the apoptotic alteration of the entire cell water and ion balance in U937 cells treated with staurosporine (STS) and etoposide (Yurinskaya et al. 2005*a*,*b*; Yurinskaya et al. 2011). However, we lacked the necessary experimental data and a proper programme code for computation of transient processes in cell ion homeostasis and analysed only apoptotic cells at a single time point, 4 h. Here, we studied ionic events during apoptosis development from 30 min to 4 h. The background data included K^+^, Na^+^, Cl^−^, and water contents and ouabain-sensitive and -resistant Rb+ influx in U937 cells that were induced to undergo apoptosis by STS. An original algorithm of the numerical solution of the cell monovalent ion flux balance equations and the programme code were developed, which allowed us to account for the continuous changes in the sodium pump activity. To our knowledge, this is the first attempt to study the dynamics of the alteration of K^+^, Na^+^, Cl^−^, and water contents during apoptosis. The approach developed to study STS-induced apoptosis in U937 cells may be recommended for identification of channels and transporters responsible for alteration of cell ion and water balance in various situations.

## Methods

### Reagents

RPMI 1640 medium and foetal bovine serum (FBS, HyClone Standard) were purchased from Biolot (Russia). STS and ouabain were from Sigma-Aldrich (Germany), Percoll was purchased from Pharmacia (Sweden). The isotope ^36^Cl^−^ was from “Isotope” (Russia). Salts were of analytical grade and were from Reachem (Russia).

### Cell cultures

U937 human histiocytic lymphoma cells were obtained from the Russian Cell Culture Collection (Institute of Cytology, Russian Academy of Sciences, cat. number 160B2). The cells were cultured in RPMI 1640 medium supplemented with 10% FBS at 37 °C and 5% CO2. For the induction of apoptosis, the cells, at a density of 1 × 10^6^ cells per ml, were exposed to STS for 0.5-4 h. All the incubations were done at 37 °C.

### Determination of cell ion and water contents

The experimental methods used in this work have been described in detail earlier (Yurinskaya et al. 2005*a*, *b*, 2011; Vereninov et al. 2007, 2008). In summary, the cells were pelleted in RPMI medium, washed five times with MgCl2 solution (96 mM) and treated with 5% trichloroacetic acid (TCA). TCA extracts were analysed for ion content. Intracellular K^+^, Na^+^ and Rb+ contents were determined by flame emission on a Perkin-Elmer AA 306 spectrophotometer. To determine the intracellular Cl^−^, the cells were cultured for 90 min or more at 37 °C in RPMI medium containing 36Cl^−^ (0.12 μCi ml^-1^). The radioactivity of ^36^Cl^−^ in TCA extracts was measured using a liquid scintillation counter (Beckman LS 6500). The intracellular Cl^−^ content was calculated, taking into account the specific activity of ^36^Cl^−^ (∼2 counts min^-1^ μmol^-1^). The TCA precipitates were dissolved in 0.1 N NaOH and analysed for protein by the Lowry procedure, with serum bovine albumin as a standard. The cell ion content was calculated in micromoles per gram of protein.

Cell water content was determined by measurements of the buoyant density of the cells in continuous Percoll gradient. Percoll solution was prepared according to the manufacturer’s instructions, and a thick cell suspension (0.1-0.2 ml, ∼ 3×10^6^ cells) was placed on the solution surface and centrifuged for 10 min at 400 × g (MPW-340 centrifuge, Poland). The buoyant density of the cells was estimated using density marker beads (Sigma-Aldrich, Germany). The water content per gram of protein, *v*_prot_, was calculated as *v*_prot_.= (1-*ρ*/*ρ*_dry_)/[0.72(*ρ*-1)], where *ρ* is the measured buoyant density of the cells and *ρ*_dry_ is the cell dry mass density, which was 1.38 g ml^-1^. The proportion of protein in dry mass was 72%.

## The sodium pump rate coefficient determination

The sodium pump rate coefficient was determined based on the assay of the ouabain-sensitive Rb+ influx and cell Na^+^ content. The cells were incubated in medium with 2.5 mM RbCl and with or without 0.1 mM ouabain for 10 min. The rate coefficient of the sodium pump (*beta*) was calculated as the ratio of the Na^+^ pump efflux to the cell Na^+^ content given the assumption of the simple linear dependence of Na^+^ efflux on cell Na^+^ in the studied range of concentrations. The pump Na^+^ efflux was calculated from ouabain-sensitive (OS) Rb+ influx assuming proportions of [Rb]o and [K]o of and 5.8 mM, respectively, and Na/K pump flux stoichiometry of 3:2.

### Calculation of the monovalent ion flux balance

The mathematical model of cell ion homeostasis and the algorithm of the numerical solution of the flux balance were described in detail earlier (Vereninov et al. 2014; 2016). The reader can reproduce all presented computed data and perform new calculations for various parameters by using the execution file to programme code BEZ01B (How to use programme code BEZ01B.zip).

This software differs from the previous BEZ01 by the additional parameter *kb*, which characterizes a decrease in the pump rate coefficient *β* with time. Symbols and definitions used are shown in **Table 1**. The input data used in calculation as file DataB.txt (see BEZ01B.zip) are the following: extracellular and intracellular concentrations (*na0*, *k0*, *cl0* and *B0*; *na*, *k* and *cl*); *kv*; the pump rate coefficient (*β*); the pump Na/K stoichiometric coefficient (*γ*); parameter *kb*; channel permeability coefficients (*pna*, *pk*, *pcl*); and the rate coefficients for the NC, KC and NKCC cotransporters (*inc*, *ikc*, *inkcc*). The results of our computations appear in the file RESB.txt (**Table 2**) after running the executable file.

**Table 1.**
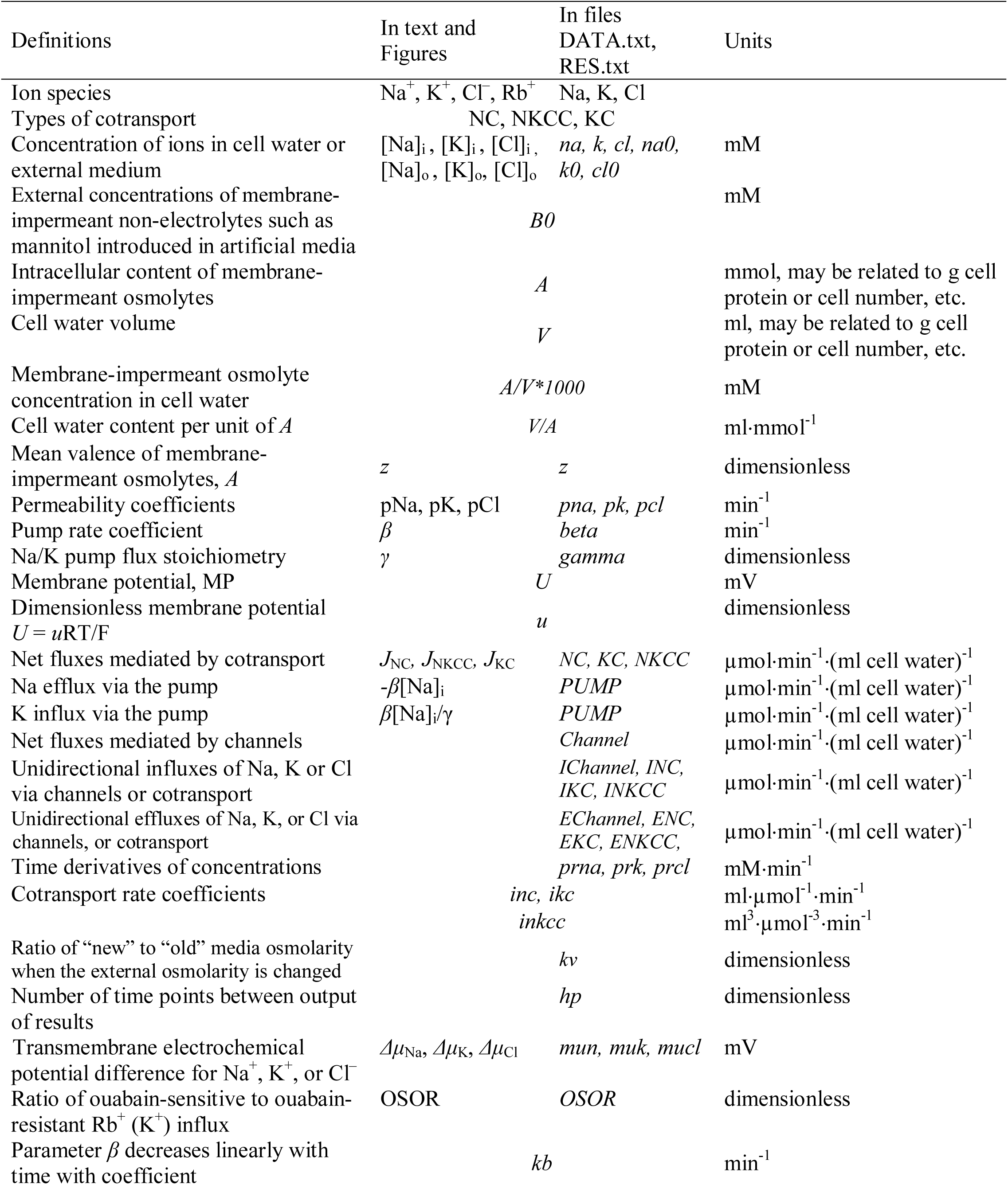
Symbols and definitions.

**Table 2.** Results of computation. Transition of the system to the balanced state as displayed in the file RESB.txt. The values similar to those for U937 cells with a rather high *U* and *mucl* were chosen for this example of a transition to the balanced state. The displayed values of fluxes as well as *OSOR* correspond to the latest time point. The values of fluxes for other moments can be obtained by setting the ecessary time interval with the *hp* value. The presented flux data do not include the fluxes involved in one-for-one exchange because they have no effect on cell ion or water content or MP and can be ignored here. The flux data clearly demonstrate how the net fluxes via different channels and transporters compensate for each other and come finally, under appropriate conditions, to a fully balanced ion distribution when the balance of influx and efflux is achieved for all ion species and *prna, prk, prcl* tend to zero.

| $t$ | $U$ | $na$ | $k$ | $cl$ | $V/A$ | $mun$ | $muk$ | $mucl$ | $prna$ | $prk$ | $prcl$ |
| --- | --- | --- | --- | --- | --- | --- | --- | --- | --- | --- | --- |
| 0 | -44.3 | 33.0 | 152.0 | 45.0 | 12.50 | -82.8 | 42.9 | 19.0 | 0.00000 | 0.00000 | 0.00000 |
| 24 | -44.5 | 36.1 | 148.9 | 45.2 | 12.52 | -80.7 | 42.2 | 19.3 | 0.07612 | -0.07696 | 0.00308 |
| ** |  |  |  |  |  |  |  |  |  |  |  |
| 216 | -44.7 | 38.0 | 147.0 | 45.1 | 12.51 | -79.5 | 41.7 | 19.4 | 0.00007 | 0.00004 | -0.00040 |
| 240 | -44.7 | 38.0 | 147.0 | 45.1 | 12.51 | -79.5 | 41.7 | 19.4 | 0.00004 | 0.00005 | -0.00032 |

\*\* Time points not shown.

(b) Parameter values (copy of the file DATAB.txt).
| $na0$ | $k0$ | $cl0$ | $B0$ | $kv$ | $beta$ | $gamma$ | $z$ | $kb$ |
| --- | --- | --- | --- | --- | --- | --- | --- | --- |
| 140 | 5.8 | 116 | 48.2 | 1.0 | 0.039 | 1.5 | -1.75 | 0 |
| $pna$ | $pk$ | $pl$ | $inc$ | $ikc$ | $inkcc$ | $A/V*1000$ | $hp$ | |
| 0.00382 | 0.022 | 0.0091 | 3E-5 | 0 | 0 | 80.0 | 240 |  |

(c) Flux balance under the balanced state.
| Net flux | PUMP | Channel | NC | KC | NKCC |
| --- | --- | --- | --- | --- | --- |
| Na | -1.4811 | 1.0451 | 0.4359 | 0.0000 | 0.0000 |
| K | 0.9874 | -0.9878 | 0.0000 | 0.0000 | 0.0000 |
| Cl | 0.0000 | -0.4363 | 0.4359 | 0.0000 | 0.0000 |
| Influx | PUMP | IChannel | INC | IKC | INKCC |
| Na | 0.0000 | 1.1012 | 0.4872 | 0.0000 | 0.0000 |
| K | 0.9874 | 0.2627 | 0.0000 | 0.0000 | 0.0000 |
| Cl | 0.0000 | 0.4082 | 0.4872 | 0.0000 | 0.0000 |
| Efflux | PUMP | EChannel | ENC | EKC | ENKCC |
| Na | -1.4811 | -0.0561 | -0.0513 | 0.0000 | 0.0000 |
| K | 0.0000 | -1.2505 | 0.0000 | 0.0000 | 0.0000 |
| Cl | 0.0000 | -0.8445 | -0.0513 | 0.0000 | 0.0000 |
| OSOR | 3.76 |  |  |  |  |

The flux equations were:s

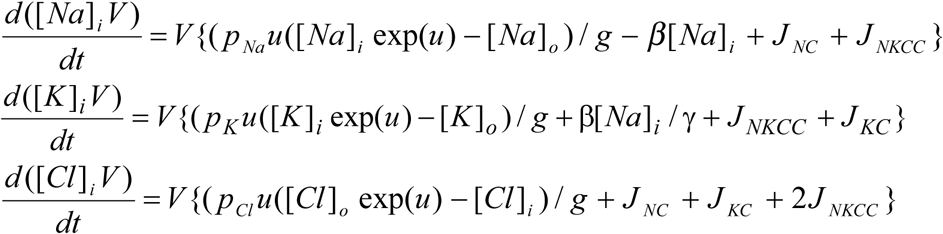

The left-hand sides of these three equations represent the rates of change of cell ion content. The right-hand sides express fluxes, where *u* is the dimensionless membrane potential related to the absolute values of membrane potential *U* (mV), as *U* = *u*RT/F = 26.7*u* for 37 °C and *g* = 1 − exp(*u*). Transmembrane electrochemical potential differences for Na^+^, K^+^, and Cl^−^ were calculated as: *Δμ*_Na_ =26.7·ln(*na/na0*)+*U*, *Δμ*_K_ =26.7·ln(*k/k0*)+*U*, and *Δμ*_Cl_ =26.7·ln(*cl/cl0*)-*U*, respectively. The values of electrochemical potential differences for Na^+^, K^+^ and Cl^−^, denoted also as *mun*, *muk* and *mucl*, are important because they show the driving force and the direction of ion movement via channels and transporters under the indicated conditions. It is the changes in *mun*, *muk* and *mucl* that are responsible for the possible fast effects of the membrane potential (MP) on ion fluxes via “electroneutral” transporters.

### Statistical analysis

Data are presented as the mean ± SEM. P < 0.05 (Student’s *t* test) was considered statistically significant.

## Results

### Computational approach to solution of the problem how the entire cell ion and water balance depends on the state of various channels and transporters

The first of the two main aims of the present study is the demonstration of the computational approach to solution of the problem how the entire cell ion and water balance depends on the parameters of various channels and transporters. The second aim is the analysis of the ion and water balance changes during apoptosis in real U937 cells. This aim is an example of using the developed approach. Some background points should be considered first. The basic mathematical model used in our approach is similar to the known model developed by pioneers for analysis of ion homeostasis in normal cells (Jakobsson, 1980; Lew & Bookchin, 1986; Lew et al. 1991). Our algorithm of the numerical solution of the flux equations and basic software were published earlier (Vereninov et al. 2014; 2016). Some minor differences in mathematical models used by previous authors consist in the number of transporters included in the calculations. Only the sodium pump and electroconductive channels were considered in the early computational studies of cell ion balance. Lew and colleagues were the first who found that the sodium pump and electroconductive channels cannot explain monovalent ion flux balance in human reticulocytes because they cannot explain the nonequilibrial Cl^−^ distribution under the balanced state without NC (Lew et al. 1991). Cotransporters NC and KC were investigated by Hernández and Cristina (1998). The NKCC cotransport was included in ion balance modelling in cardiomyocytes (Terashima et al. 2006). Our software accounts for Na^+^, K^+^, and Cl^−^ channels, the sodium pump and the NC, KC and NKCC cotransporters. We found that NC is necessary as a rule in the calculation of the resting monovalent ion flux balance in U937 cells, while NKCC and KC are not. Nevertheless, the parameters characterizing these two transporters are present in our code, and fluxes via transporters can be accounted for if these parameters differ from zero.

Two points may worry experimentalists. First, the sodium pump activity is characterized by a single rate coefficient. However, a set of ion binding sites are known in the pump, and its operation kinetics in biochemical studies is described commonly by more than one parameter. The single rate coefficient is used because the evaluation of the properties of all the ion binding sites of the pump in experiments in whole cells is infeasible and because it appears to be quite sufficient for the calculation of entire-cell ion homeostasis. This idea was demonstrated by the quantitative prediction of the dynamics of monovalent ion redistribution after stopping the sodium pump (Vereninov et al. 2014, 2016). Single rate coefficients for characterizing the ion carriage kinetics via transporters are commonly used for the same reason. The second point causing disapproval might be that an integral permeability coefficient is used in the calculation of the flux balance for all Na^+^ or K^+^ or Cl^−^ channels, whereas a great variety of channels for each ion species is located in the plasma membrane. The single permeability coefficients are commonly used in analysis of the entire-cell flux balance because in an analysis of such a complex system with many channels and transporters, the matter of primary importance is to understand whether ion flux changes due to alteration of the force driving the ions or by properties of the channel per se.

### Computation of ion flux balance in cells similar to U937 cells

Parameters in absolute units are used in our calculations. Their initial, “standard”, values are obtained from the calculation based on the distribution of monovalent ions and ouabain-sensitive Rb+(K^+^) influx measured in cells under normal physiological conditions and the cell balanced state. The parameters varied until the tested values give a calculated entire ion and water homeostasis similar to that in real cells. The “standard” parameters can vary in real cells depending on the cell physiological state, the age of the culture, the conditions of cell cultivation, etc.

Nevertheless, these parameters can remain invariant under a varying environment. We found that the kinetics of the disturbance of cell ion and water balance caused by blocking the sodium pump when the intracellular K^+^/Na+ ratio is highly changed and even reversed can be predicted sufficiently well by calculation with the invariant parameters (Vereninov et al. 2014; 2016). A set of examples is presented in **Figure 1** to show how changes in a single channel or transporter species (one permeability coefficient or rate constant) can alter the intracellular concentrations of all major ions, cell water content and the MP. Unlike pNa and *inc*, changes in pK or pCl lead, over the course of 60-100 min, to a new balanced state. Even this small set of examples demonstrates that some effects seem to be unexpected at first sight. Intracellular K^+^ concentration decreases monotonically with the pNa increase, while the intracellular K^+^ content decreases initially and increases further due to superposition of the initial drop MP and the slow increase in cell water-volume. An increase in the coupled equivalent transport Na^+^ and Cl^−^ (*inc*) causes a decrease in cell K^+^ concentration, [K^+^], and, in contrast, an increase in cell K^+^ content because of changes in cell water volume and in MP. The [K^+^] and MP are shifted in this case in opposite directions. It should be stressed that the effects of parameter variation are highly dependent on the cell species. Our previous paper presented the typical dependences for the cells with high MP and high intracellular K/Na ratio, for the cells with low MP and high K/Na ratio (high potassium erythrocytes) and for the low-MP and low-K/Na-ratio cells (low-potassium erythrocytes of some carnivores and ruminants) (Vereninov et al., 2014). Cells such as U937 and their variant with relatively high *mucl* (19.4 mV) are chosen as an example in **Figure 1** to make the pCl effect more obvious.

**Figure 1.**
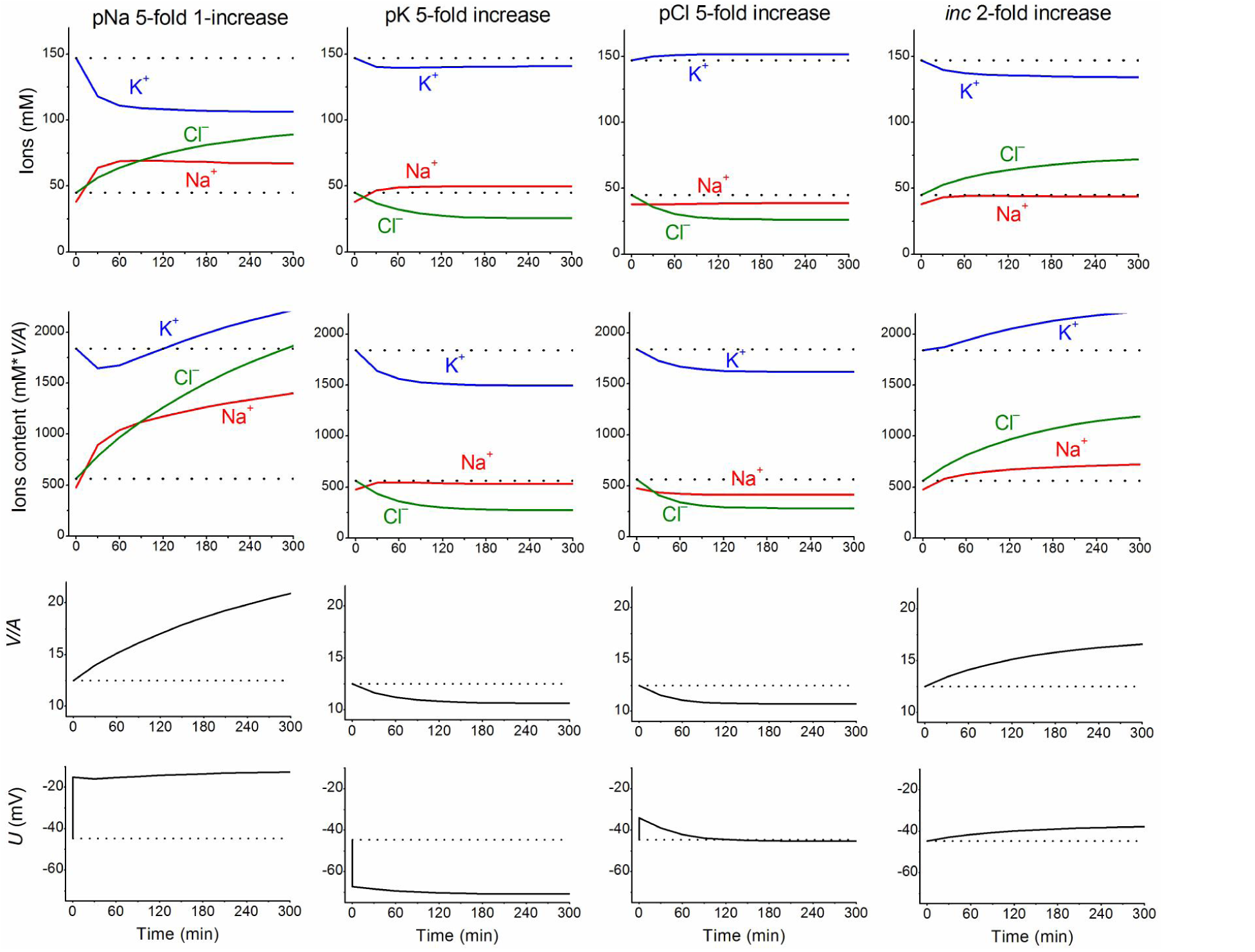
**The calculated effects of an abrupt increase in the permeability coefficients of K^+^, Na^+^, Cl^−^ channels, or the NC cotransport rate coefficient on cell K^+^, Na^+^, and Cl^−^ content and concentrations, water-volume (*V/A*) and MP (*U*)**. The data were calculated by using the software BEZ01B. The initial parameters were *na0* 140, *k0* 5.8, *cl0* 116, *B0* 48.2, *kv* 1, *beta* 0.039, *gamma* 1.5, *pna,* 0.00382, *pk* 0.022, *pcl* 0.0091, *inc* 0.00003, *ikc* = *inkcc* = 0, *kb* 0, and *hp* 300, i.e., much like U937 cells; the changed parameters are shown on the plots.

The ion and water redistribution caused in U937 cells by stopping the sodium pump was studied earlier in silico and in an experiment (Vereninov et al. 2014; 2016). Here, it is interesting to demonstrate this case as an example of asynchrony in changes of K^+^, Na^+^ and Cl^−^ after blocking the pump (**Figure 2**). In the earlier stage, the electroneutrality of the net ion fluxes is achieved mainly by the balance of fluxes K^+^ outward and Na^+^ inward via channels, whereas, later, the Cl^−^ influx becomes significant. There is no alteration of total intracellular osmolytes during the equal K^+^/Na+ exchange, and it is for this reason that no swelling occurs after blocking the pump for a rather long time. It should be stressed that no specific carrier is responsible for the balanced K^+^/Na+ exchange. This result is realized via electroconductive channels only due to the dependence of fluxes on MP. The long-term balanced state in monovalent ions and water distribution after stopping the pump is unattainable, and cell swelling will go on infinitely. However, the kinetics of the entire process may be different in dependence on the *pcl* level. It should be noted that cell water content and intracellular concentration are changing synchronously. It is the low Cl^−^ channel permeability that saves real cells from swelling for a long time after blocking the sodium pump.

**Figure 2.**
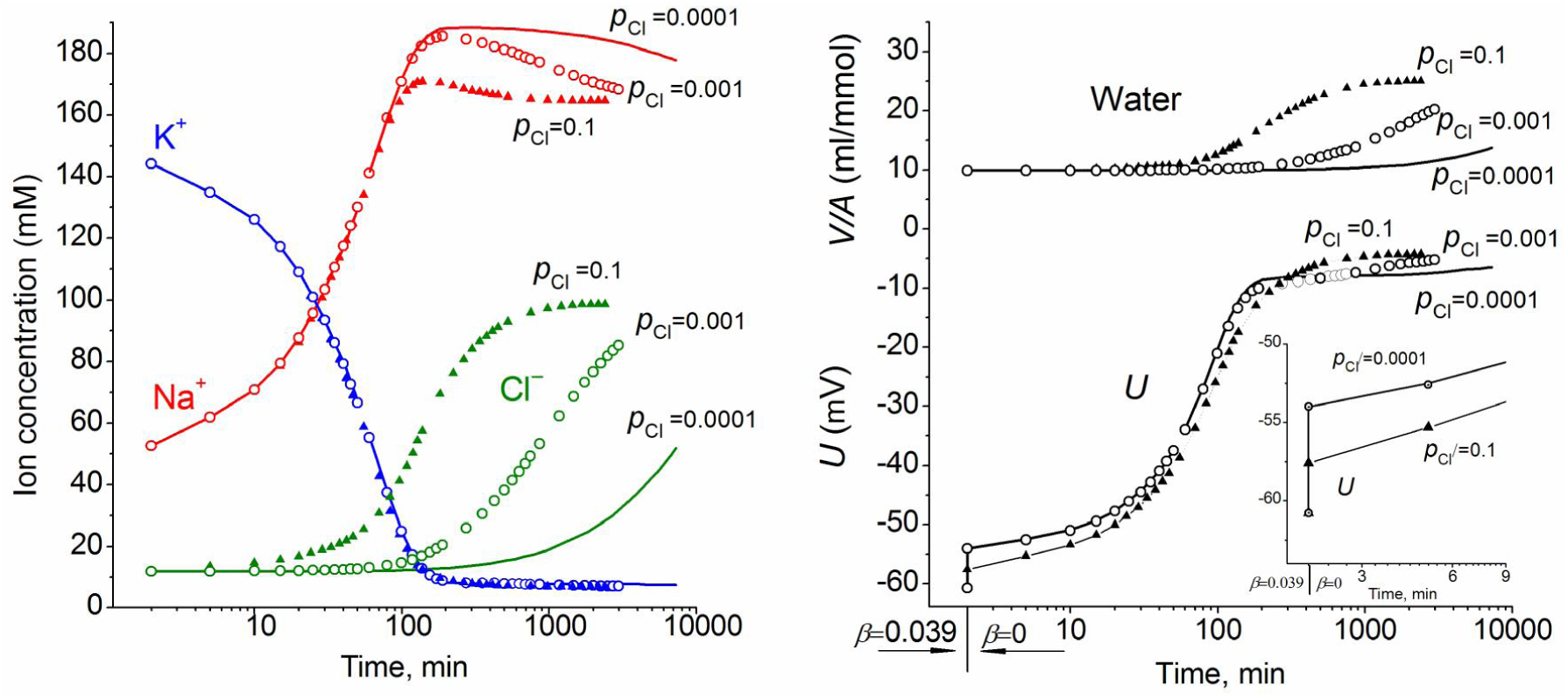
The effect of pCl on the time course of the ion and water balance disturbance caused by turning off the pump. The data were calculated by using the software BEZ01B with the following parameters: *na0* 140, *k0* 5.8, *cl0* 116, *B0* 48.2, *kv* 1, *na* 52.6, *k* 144.2, *cl* 11.9, *beta* 0 (at the initial balanced value of 0.039), *gamma* 1.5, *pna* 0.006, *pk* 0.06, *pcl* 0.1 (triangles) or 0.001 (circles) or 0.0001 (solid lines), *inc* = *ikc* = *inkcc* = 0, *kb* 0.

### Changes in K^+^, Na^+^, Cl^−^ and water contents during early apoptosis in U937 cells induced by STS

Most data on the redistribution of monovalent ions during apoptosis relates to the 4-5 h stage (see references in Arrebola et al. 2005*a*). Our simultaneous determination of K^+^, Na^+^, Cl^−^ and water contents in U937 cells treated with STS for 4 h was published earlier (Yurinskaya et al. 2011). The data related to this apoptosis stage confirmed the osmotic mechanism of AVD, i.e., they showed that the water loss was caused mostly by the loss of the total monovalent ion content and much less by a decrease in content of the “impermeant intracellular anions”, *A*^−^. The initial changes in all major monovalent ions and water content during apoptosis have been studied much less, although the early cell shrinkage is supposed to be crucial for triggering apoptosis. Our current data on the changes in ion and water content during STS apoptosis in U937 cells with the earliest time point 30 min are presented in **Table 3**. The values of the independently determined ion content and water content correspond to the osmotic mechanism of AVD at the early stages as well as at the 4 h stage studied before. A decrease in cell water fits a decrease in the total content of intracellular osmolytes. The data on water content in Table 3 were obtained by the best method, i.e., by cell buoyant density. These data agree well with the data obtained by using a Coulter counter and flow cytometer (Yurinskaya et al. 2017). Calculation of the changes in K^+^, Na^+^, and Cl^−^ net fluxes underlying the changes in cell ion and water content shows that for the first hour, the K^+^ loss is electrically balanced predominantly by the Cl^−^ loss, whereas later it is mostly balanced by the Na^+^ gain (Table 4, last columns).

**Table 3.** Changes in K^+^, Na^+^, Cl^−^ and water contents during the early stages of STS apoptosis in U937 cells. Means ± SEM from 3 independent experiments with duplicate determinations.

| Time | $K^+$ | $Na^+$ | $Cl^-$ | $A^-$ | Water |
| --- | --- | --- | --- | --- | --- |
| min | $\mu\text{Eqv}\cdot(\text{g prot.})^{-1}$ | | | | ml/g |
| 0 | $712 \pm 22$ | $192 \pm 8$ | $246 \pm 11$ | 658 | $6.08 \pm 0.08$ |
| 30 | $615 \pm 12$ | $175 \pm 10$ | $133 \pm 4$ | 657 | $5.37 \pm 0.21$ |
| 120 | $595 \pm 13$ | $179 \pm 4$ | $109 \pm 5$ | 665 | $4.70 \pm 0.05$ |
| 240 | $493 \pm 21$ | $261 \pm 5$ | $117 \pm 4$ | 637 | $4.85 \pm 0.08$ |

**Table 4.** Changes in cell [K^+^], [Na^+^], [Cl^−^], MP (U) and net fluxes calculated upon a linear decrease in the pump rate coefficient and stepwise changes in the K^+^, Na^+^ and Cl^−^ channel permeability corresponding to the experimental data on apoptotic alteration of K^+^, Na^+^, and Cl^−^ concentrations and pump fluxes. The initial parameters were *na0* 140, *k0* 5.8, *cl0* 116, *B0* 48.2, *kv* 1, *na* 32, *k* 117, *cl* 40, *gamma* 1.5, *inc* 0.000003, and *kb* 0.000068. The changed parameters, including *beta, pna*, *pk*, and *pcl*, are shown in the Table. ** Time points not shown. The data were obtained by using the code BEZ01B. The time of channel alteration is indicated by horizontal lines. Outward net fluxes are defined as negative.

| Time,<br>min | $\beta$ | $p_k$ | $p_{na}$ | $p_{cl}$ | $U$ | $na$ | $k$ | $cl$ | $V/A$ | $\mu_{un}$ | $\mu_k$ | $\mu_{cl}$ | Net fluxes | | |
| --- | --- | --- | --- | --- | --- | --- | --- | --- | --- | --- | --- | --- | --- | --- | --- |
| | | | | | | | | | | | | | $Na^+$ | $K^+$ | $Cl^-$ |
| Before | 0.029 | 0.0115 | 0.0041 | 0.0125 | -29.9 | 32 | 117 | 40 | 8.26 | -69.3 | 50.3 | 1.5 | 0 | 0 | 0 |
| 0 | 0.029 | 0.03 | 0.003 | 0.068 | -34.6 | 32.0 | 117 | 40.0 | 8.26 | -74.0 | 45.6 | 6.2 | 0 | 0 | 0 |
| 10 | 0.028 |  |  |  | -38.5 | 32.3 | 116.4 | 33.5 | 7.82 | -77.7 | 41.6 | 5.3 | -0.119 | -0.622 | -0.733 |
| 20 | 0.028 |  |  |  | -42.0 | 32.7 | 115.7 | 28.4 | 7.51 | -80.8 | 37.9 | 4.4 | -0.065 | -0.481 | -0.540 |
| 30 | 0.027 |  |  |  | -45.0 | 33.3 | 114.9 | 24.6 | 7.29 | -83.3 | 34.8 | 3.5 | -0.020 | -0.366 | -0.381 |
| 60 | 0.025 | 0.02 | 0.003 | 0.068 | -44.8 | 33.8 | 114.3 | 22.3 | 7.16 | -82.8 | 34.8 | 0.8 | 0.031 | -0.079 | -0.048 |
| 120 | 0.021 |  |  |  | -45.1 | 37.7 | 110.3 | 21.7 | 7.13 | -80.1 | 33.5 | 0.3 | 0.086 | -0.078 | 0.008 |
| ** |  |  |  |  | ..... |  |  |  |  |  |  |  |  |  |  |
| 210 | 0.015 | 0.02 | 0.003 | 0.068 | -43.2 | 47.2 | 100.9 | 23.2 | 7.21 | -72.2 | 33.1 | 0.1 | 0.139 | -0.110 | 0.029 |
| 240 | 0.013 |  |  |  | -42.2 | 51.4 | 96.7 | 24.0 | 7.25 | -68.9 | 33.0 | 0.1 | 0.163 | -0.127 | 0.036 |

The apoptotic changes in ion content obtained in our study by flame photometry and radiotracer assay are very close to the data obtained by the X-ray microanalysis in U937 cells during several types of apoptosis, including early STS apoptosis (Arrebola et al. 2005*b*, 2006). Unlike that X-ray microanalysis study, we could more easily validate changes in cell water content during apoptosis and therefore better estimate ion concentrations per cell water volume. This approach enabled us to calculate the entire cell electrochemical model and, in this way, to identify channels and transporters critical for AVD.

### Matching the real and calculated changes in cell K^+^, Na^+^ and Cl^−^ concentrations during apoptosis

The real changes in Na^+^, K^+^, Cl^−^ and water contents during STS apoptosis in U937 cells differ from the calculated example presented in **Figure 1**. Evidently, other changes in rate parameters during the transient process can occur in real cells. Indeed, a decrease in the sodium pump activity is a peculiar feature of apoptosis and has been revealed without any computation, particularly in U937 cells treated with STS (Arrebola et al. 2005*a, b*; Vereninov et al. 2008; Yurinskaya et al. 2010, 2011). The rate coefficient of the sodium pump can be validated by OS Rb+ influx and intracellular Na^+^ content (Vereninov et al. 2014, 2016). We found that the OS Rb+ influx for the first 4 h of STS apoptosis in U937 cells decreased mostly linearly (Yurinskaya et al. 2010).

The linear decrease in the sodium pump rate coefficient with time was accounted for in the current programme code BEZ01B. **Figure 3A** shows the transient process during STS apoptosis in U937 cells calculated under the assumption that the sodium pump rate coefficient decreases linearly due to the decrease in the coefficient *kb*, as found by OS-Rb+ influx assay in experiment. The values calculated for this simplest model (lines) correspond approximately to the real ion concentrations (symbols) for K^+^ (circles) and Na^+^ (triangles) but differ significantly for Cl^−^ (squares). The additional assumption that triggering apoptosis is accompanied by stepwise increases in *pcl* and *pk* and by a slight decrease in *pn*a improves the agreement between calculated and real values for Cl^−^ in the first 30-40 min, but not later **(Figure 3B**). A change in pCl alone is ineffective because of the small *Δμ*Cl. The agreement may be obtained for the whole 4 h time interval by assuming that *pk* further decreases (**Figure 3C**). How unique is the found fitting? By trial and error, we found that a *pna* decrease and a *pk* increase alone without a *pcl* increase could be sufficient to get agreement between real and calculated chloride concentrations for the first 30 min. However, this case should be rejected because the value OSOR becomes unacceptably low.

**Figure 3.**
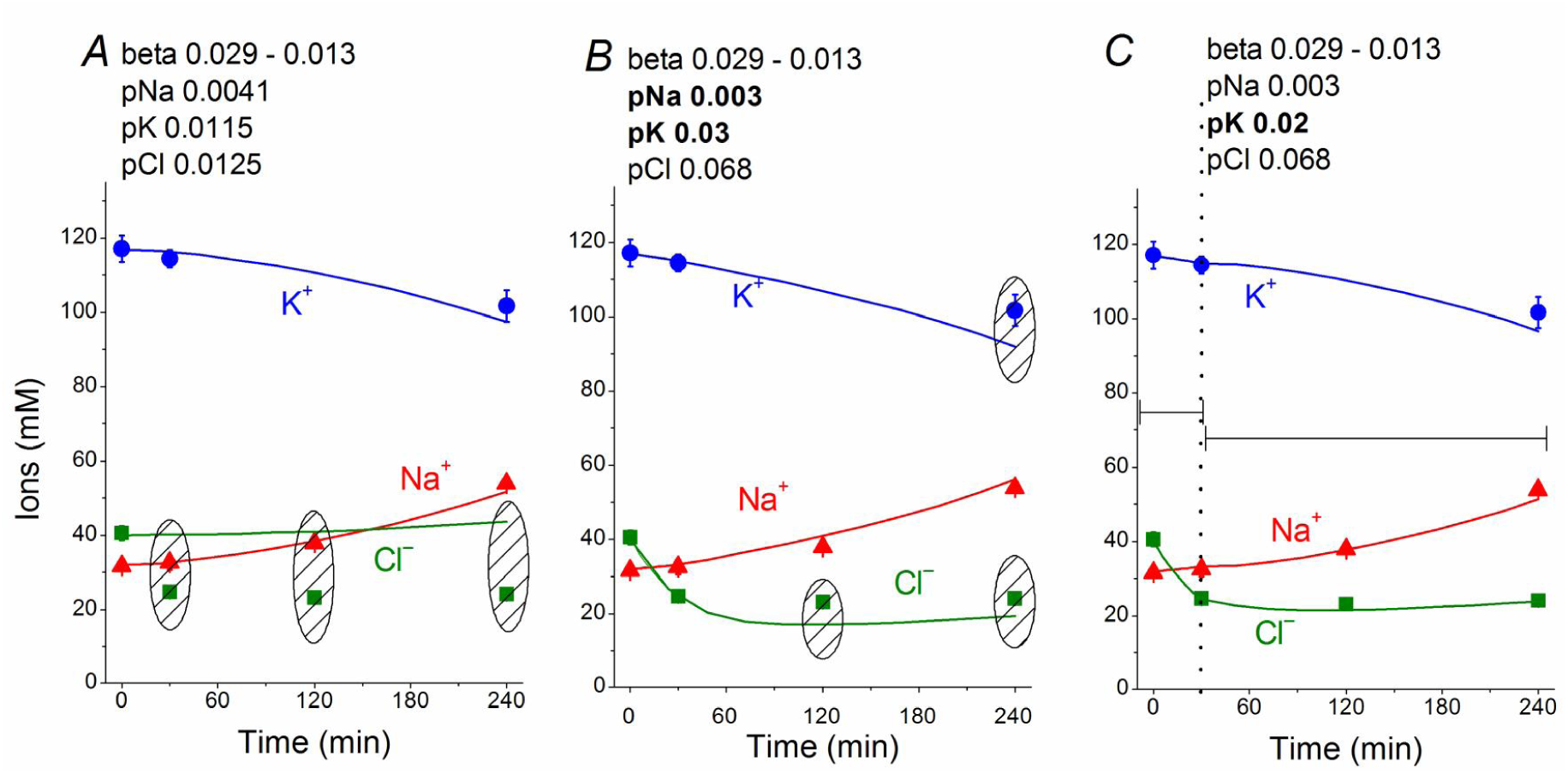
Time course of [K^+^], [Na^+^] and [Cl^−^] in real U937 cells treated with 1 µM STS (symbols) and calculated (lines) for different parameter datasets. Symbols – experimental data, means ± SEM from 3 independent experiments with duplicate determinations. Small SEM values are masked by symbols. Lines – calculated data obtained for the parameters indicated on the graphs. The initial parameters were *na0* 140, *k0* 5.8, *cl0* 116, *B0* 48.2, *kv* 1, *na* 32, *k* 117, *cl* 40, *beta* 0.029, *gamma* 1.5, *pna* 0.0041, *pk* 0.0115, *pcl* 0.0125, *inc* 0.000003, *ikc* = *inkcc* = 0, and *kb* 0.000068. The changed parameters are shown in the layers head. (A) Linear decrease in *beta* only. (B) Ddecrease in *beta* and changes in *pna, pk,* and *pcl*. (C) Additional decrease in *pk*. Shaded regions show significant disagreement of experimental and predicted values. The calculated data were obtained by using code BEZ01B.

The joint effect of pK, pNa and pCl shift is interesting. The cells shown in **Figure 3** had initially a rather low *U* (–29.9 mV) and a *mucl* of approximately 1.5 mV under the normal state (**Table 4**). The variation of pCl alone at so small a *mucl* has no significant effect. A decrease in pNa hyperpolarizes cells promptly, and an increase in pK alone hyperpolarizes cells as well (**Table 5**).

**Table 5.** The effects of pK, pNa and pCl shifts at the initial stage of apoptosis on *U*, *mucl* and ion concentrations. The initial parameters were as follows: *na0* 140, *k0* 5.8, *cl0* 116, *B0* 48.2, *kv* 1, *beta* 0.029, *na* 32, *k* 117, *cl* 40, *gamma* 1.5, *inc* 0.000003, *kb* 0.

| *t* | *pk* | *pna* | *pcl* | *U* | *na* | *k* | *cl* | V/A | *mun* | *muk* | *mucl* |
| --- | --- | --- | --- | --- | --- | --- | --- | --- | --- | --- | --- |
| pK shift |  |  |  |  |  |  |  |  |  |  |  |
| initial | 0.0115 | 0.0041 | 0.0125 | -29.9 | 32.0 | 117.0 | 40.0 | 8.26 | -69.3 | 50.3 | 1.5 |
| 0 | 0.03 | 0.0041 | 0.0125 | -41.1 | 32.0 | 117.0 | 40.0 | 8.26 | -80.6 | 39.1 | 12.7 |
| 15 |  |  |  | -42.1 | 35.1 | 113.7 | 36.6 | 8.03 | -79.0 | 37.4 | 11.3 |
| 30 |  |  |  | -42.9 | 37.2 | 111.5 | 33.8 | 7.83 | -78.3 | 36.0 | 10.0 |
| 45 |  |  |  | -43.6 | 38.6 | 110.0 | 31.5 | 7.69 | -78.0 | 34.9 | 8.8 |
| 60 |  |  |  | -44.2 | 39.2 | 109.0 | 29.5 | 7.57 | -78.0 | 34.1 | 7.7 |
| pNa shift |  |  |  |  |  |  |  |  |  |  |  |
| initial | 0.0115 | 0.0041 | 0.0125 | -29.9 | 32.0 | 117.0 | 40.0 | 8.26 | -69.3 | 50.3 | 1.5 |
| 0 | 0.0115 | 0.003 | 0.0125 | -36.6 | 32.0 | 117.0 | 40.0 | 8.26 | -76.0 | 43.6 | 8.2 |
| 15 |  |  |  | -37.2 | 30.6 | 118.3 | 38.0 | 8.12 | -77.8 | 43.3 | 7.4 |
| 30 |  |  |  | -37.8 | 29.7 | 119.1 | 36.2 | 8.00 | -79.2 | 42.8 | 6.7 |
| 45 |  |  |  | -38.4 | 29.2 | 119.5 | 34.6 | 7.90 | -80.2 | 42.4 | 6.1 |
| 60 |  |  |  | -38.9 | 28.9 | 119.7 | 33.3 | 7.81 | -81.0 | 41.9 | 5.6 |
| pK and pNa shift |  |  |  |  |  |  |  |  |  |  |  |
| initial | 0.0115 | 0.0041 | 0.0125 | -29.9 | 32.0 | 117.0 | 40.0 | 8.26 | -69.3 | 50.3 | 1.5 |
| 0 | 0.03 | 0.003 | 0.0125 | -44.8 | 32.0 | 117.0 | 40.0 | 8.26 | -84.2 | 35.4 | 16.4 |
| 15 |  |  |  | -46.3 | 32.5 | 116.3 | 35.6 | 7.96 | -85.3 | 33.8 | 14.7 |
| 30 |  |  |  | -47.6 | 32.8 | 115.8 | 31.9 | 7.72 | -86.4 | 32.3 | 13.1 |
| 45 |  |  |  | -48.8 | 33.1 | 115.4 | 28.8 | 7.53 | -87.3 | 31.1 | 11.5 |
| 60 |  |  |  | -49.7 | 33.3 | 115.0 | 26.2 | 7.38 | -88.1 | 30.0 | 10.0 |
| pK, pNa and pCl shift |  |  |  |  |  |  |  |  |  |  |  |
| initial | 0.0115 | 0.0041 | 0.0125 | -29.9 | 32.0 | 117.0 | 40.0 | 8.26 | -69.3 | 50.3 | 1.5 |
| 0 | 0.03 | 0.003 | 0.068 | -34.6 | 32.0 | 117.0 | 40.0 | 8.26 | -74.0 | 45.6 | 6.2 |
| 15 |  |  |  | -40.4 | 32.3 | 116.2 | 30.7 | 7.65 | -79.6 | 39.7 | 4.9 |
| 30 |  |  |  | -45.2 | 32.6 | 115.6 | 24.4 | 7.28 | -84.2 | 34.7 | 3.6 |
| 45 |  |  |  | -48.7 | 32.9 | 115.1 | 20.5 | 7.07 | -87.4 | 31.1 | 2.5 |
| 60 |  |  |  | -50.9 | 33.2 | 114.7 | 18.3 | 6.95 | -89.4 | 28.8 | 1.6 |

As a result, the pCl increase becomes effective and sufficient to get both the necessary agreement between real and calculated chloride concentrations for the initial 30-40 min and the necessary OSOR.

The *inc* and pCl parameters change cell water and [Cl]_i_ in opposite directions (**Figure 1**). However, we could not replace the pCl increase with the *inc* decrease in our fitting procedure, as the [Cl]_i_ decrease in the latter case is too small. We came to conclusion that an increase in pCl is a critical factor for the complex water and ion rearrangement at the initial stage of STS apoptosis in U937 cells, whereas its role becomes less significant or even non-significant later.

We conclude that the redistribution of K^+^, Na^+^, and Cl^−^ underlying AVD in the studied U937 cells treated with STS is caused (1) by a progressive linear decrease in the sodium pump rate coefficient from the initial 0.029 to 0.013 at 4 h, (2) by a significant increase in pCl (0.0125 to 0.068) and changes in pK (0.0115 to 0.03 and later 0.02), and (3) by a moderate decrease in pNa (0.0041 to 0.003). The most critical factors for changes in cell K^+^ and Na^+^ are suppression of the pump, an increase in pK and a decrease in pNa, whereas the early decreases in Cl^−^ and water content (early AVD) are associated primarily with an increase in pCl by approximately 5 times and an increase in pK by approximately 2.6 times.

The calculated MP in the considered model of apoptosis was slightly hyperpolarizing (by 12 mV). Our preliminary results from flow cytometry using DiBAC_4_(3) did not show a significant change in MP under STS-induced apoptosis (unpublished data). These results differ from previous reports of MP depolarization during apoptosis, e.g., in the Fas-L induced apoptosis of Jurkat cells (Franco et al. 2006). Further studies are required to determine whether MP changes are highly dependent on the apoptosis inducer and/or on the cell species or whether cell depolarization occurs in more severe apoptosis.

## Discussion

Monovalent ion channels and transporters are involved in apoptosis (Burg et al. 2006; Lang et al. 2007; Dezaki et al. 2012; Orlov et al. 2013; Hoffman et al. 2009, 2015; Kondratskyi et al. 2015; Wanitchakool et al. 2016; Lang & Hoffmann 2012; Pedersen et al. 2016; Jentsch, 2016). However, this phenomenon may be caused by two reasons: because the monovalent ions are the major cell volume regulators and should be responsible for AVD only for this reason, or because they also play important roles in cell signalling by affecting MP. It is not easy to distinguish these two causes at present. We aimed to answer the question how alteration of distinct channels and transporters affects the balance of monovalent ion fluxes across the cell membrane, cell water content and MP at apoptosis. We studied the time course of the monovalent ion balance redistribution during the first 4 h development of apoptosis induced in U937 cells treated with STS as the established model of apoptosis with significant AVD. Apoptosis in U937 cells is accompanied by rapid changes in light scattering and cell water (volume) balance, whereas the positive annexin test and intensive generation of apoptotic bodies are revealed, starting at 3-4 h (Yurinskaya et al. 2017). The identification of channels and transporters responsible for the observed changes in monovalent ion distribution, water balance and the sodium pump fluxes was based on the computational modelling of these changes. Such an approach was applied here to study apoptosis for the first time, although the monovalent flux balance under the normal physiological state and during redistribution of ions due to stopping the sodium pump has been calculated successfully before (see ref. in: Vereninov et al. 2014, 2016). Our previous code was modified currently to account for a continuous decrease in the sodium pump rate coefficient.

One of the most detailed studies of the kinetics of the monovalent ion balance rearrangement during apoptosis was performed by X-ray microanalysis and in U937 cells treated with STS in particular (Arrebola et al. 2005*a, b*). The experimental data obtained by flame emission and radiotracer assays in our study agree very well with the data obtained by this quite different method. Unfortunately, the accurate cell water content evaluation is hard to combine with the X-ray elemental microanalysis. Therefore, the complete mathematical model of the monovalent ion flux balance could not be developed using only those data.

Earlier we tried to relate the changes in ion and water contents to the monovalent fluxes in the sodium pump, K^+^, Na^+^, and Cl^−^ channels and certain cotransporters in U937 cells after 4 h of STS-induced apoptosis (Yurinskaya et al. 2011). We came to the conclusion that the sodium pump suppression accompanied by a decrease in Na^+^ channel permeability might be responsible for AVD under the considered conditions. However, our current computational tool had not been developed at that time, the experimental data were limited to single time point 4 h, and the assumption was used that the balanced monovalent ion distribution is reached at 4 h of STS-induced apoptosis in U937 cells. More complete current data show that the cells at the 4 h time point are far from the balanced state (Table 3). There is significant Na^+^ gain (0.163) that is by ¾ balanced by K^+^ leak (0.127) and ¼ by the gain of Cl^−^ (0.036).

Currently, we substantially revised and developed our previous conception of the participation of the major channels and transporters in AVD during the STS-caused apoptosis of U937 cells. It remains true that a slow decrease in the sodium pump activity is a primary factor responsible for AVD at the late (4 h) stage of apoptosis. Recalculation of the data published earlier (Yurinskaya et al. 2011) with use of the current programme code and without intricate hypotheses confirmed a decrease in pNa at 4 h. The current data show that the pNa decrease at 4 h is significant indeed. The most interesting and important phenomenon is a more than 5-fold increase in the Cl^−^ channel permeability, which is much more important at the early stage. It is remarkable that the effect of the pCl increase disappears further because of a decrease in intracellular Cl^−^ concentration and associated decrease in chloride electrochemical potential difference, *Δμ*Cl (*mucl* in Table 4). The effects of the early increase in pK and a decrease in pNa on *U* are significant because they lead to an increase in *Δμ*Cl that drives chloride outward. A large body of electrophysiological evidence published recently indicates that the state of the chloride channels can change upon initiation apoptosis (Hoffmann et al. 2015; Kondratskyi et al. 2015; Wanitchakool et al. 2016; Pedersen et al. 2016; Jentsch, 2016). However, there were no attempts to use these data for the quantitative description of early AVD.

The current computations show that changes in not a single type of channel but in K^+^, Na^+^, and Cl^−^ channels and in the sodium pump are responsible for the apoptotic ion balance alteration and that the effect of various channels and transporters on ion balance may be different at different stages of apoptosis. Certainly, the question arises how many parameters can provide an accordance between the calculated and real data? The computation enables us to answer this question, although certain time may be needed. In the case of STS-induced apoptosis in U937 cells in our experiments, we can exclude alternative variants by taking into account additionally the value of OSOR, which appeared to be different in different parameter setups, giving sufficiently good accordance between the real and calculated data. In other cases, the problem could be solved probably not by using OSOR but by some other way. The computation shows also how the real behaviour of cells should depend on the initial state of cells. Certainly, as soon as basic experimental data vary, the obtained numerical values of parameters will vary also. We can see from our long experience studying ion balance in cultured cells that the variability of the cell physiological state rather than the inaccuracy of assays hamper the quantitative description of cell ion and water balance.

A skeptical view is spread among the experimentalists on the using calculations in analysis of the ion flux balance in cells. There is also a great deal of sometimes convoluted discussion about the merits and validity of certain assumptions that need to be made for the models and real data to be reconciled. As believed it is very difficult to accurately assess the value of the models and their conclusions. In this regard, we should note the following. If a required set of experimental data is a unique solution appears independently on any hypotheses on the number and types of channels and transporters which could present in cell membrane. Our system of the flux equations accounts all currently known types of ion transfer across membrane characterized only by the ion driving forces: electrochemical potential difference for movement of single ion species (electrodiffusion through electroconductive channels), the sum of electrochemical potential differences for the linked movement of several species of ions (cotransport, countertransport), and a combination of the electrochemical and chemical potential differences in case of the Na,K-ATPase pump. Computer decides what number of transporting units of each type should be for implementation of two physically mandatory demands and which ion pathways do not play a role under given ion conditions. The mandatory demands are electroneutrality of the any macroscopic ion redistribution and osmotic balance between distensible animal cell and the medium. Any hypothesis on the mechanism regulating cell water and ion content or membrane potential must be checked for these demands implementation. This cannot be done without computation in system with a numerous species of ions and a numerous ion pathways. Experimentalists avoid calculation and prefer using inhibitors and genetic cell modification simply because there is no sufficiently suitable tool for computation. We attempted to reduce computational tool deficiency.

## Conclusions

The experimental data on the time course of K^+^, Na^+^, and Cl^−^ concentrations and ouabain-sensitive and -resistant Rb+ influx in U937 cells treated with STS for 0.5-4 h enabled us to evaluate the changes in the pump rate coefficient and to compute alterations of the K^+^, Na^+^, and Cl^−^ channel permeability coefficients associated with the initial stages of apoptosis and AVD.

The redistribution of K^+^, Na^+^, and Cl^−^ underlying AVD in U937 cells is caused (1) by a progressive decrease in the sodium pump rate coefficient from an initial 0.029 to 0.013 at 4 h, (2) by a significant increase in pCl (0.013 to 0.068) and increases in pK (0.012 to 0.03, later 0.02), and (3) by a moderate decrease in pNa (0.004 to 0.003). The most critical factors for changes in cell K^+^ and Na^+^ are the suppression of the pump, an increase in pK and a decrease in pNa, whereas the early decrease in Cl^−^ and water content (early AVD) are associated primarily with an increase in pCl by approximately 5 times and an increase in pK by approximately 2.6 times.

Our approach demonstrates how to calculate the dependence of cell ion and water balance on the states of channels and transporters in the plasma membrane.

## Supporting information

## Acknowledgments

We thank Dr. Tatyana Goryachaya for excellent assistance in the experiments with cells. The research was supported (VEY, AAV) by the Grant of Russian Federation (No. 0124-2018-0003).

## Author contributions

All authors contributed to the design of the experiments, performed the experiments, and analysed the data. A.V. wrote the manuscript with input from all authors. All authors have approved the final version of the manuscript and agree to be accountable for all aspects of the work. All persons designated as authors qualify for authorship, and all those who qualify for authorship are listed.

## Supporting information

Executional file to the programme code BEZ01B and Instruction: How to use programme code BEZ01B.zip. This file is attached to the article electronic version.

